# SatCHM (Satellite Canopy Height Model): Leveraging deep learning for site-specific sub-meter canopy height predictions

**DOI:** 10.64898/2026.08.25.728853

**Authors:** Mia Mitchell, Charles Abolt, Zachary Crennen, Agnese Marcato, Adam Atchley

## Abstract

High-resolution monitoring of forest structure and productivity is essential for effective natural resource management. However, monitoring approaches such as field-based forest inventories or extensive lidar campaigns are costly, time-intensive, and spatially limited. Therefore, inexpensive and accessible methods are needed. SatCHM (Satellite Canopy Height Model) was developed to be an accessible and open-source tool for researchers, allowing for site-specific and temporally flexible predictions of canopy height with limited computational resources. SatCHM requires four inputs: panchromatic satellite imagery, solar and sensor angle metadata of satellite imagery, digital elevation models (DEMs), and lidar-produced CHMs for an area of interest. After SatCHM pre-processes inputs, data is loaded into a collection of convolutional neural networks (CNNs) for image-to-image regression. This ensemble cooperates to yield high-resolution predictions (up to 0.5-meter) of three-dimensional tree structure with discernible tree crowns across a broader defined area of interest. After calculating the mean absolute error for each prediction output, the median of these mean absolute errors was 6.06 meters.

## Introduction

Canopy height models (CHMs) are useful tools for understanding forest ecosystem responses under anthropogenic global change [1, 2]. Changes in biomass are interlinked with global and local carbon cycles [3, 4]. Therefore, scientists require tools to accurately measure vegetation throughout time and space [5, 6]. High-resolution CHMs are historically produced from air-borne lidar [7]. Although the use of lidar for forest applications is increasing, high surveying costs still constrain the ability to carry out large-scale repeated campaigns [8].

Recent advances in machine learning enable CHM generation from satellite imagery [9–11]. Satellite imagery can be used to fill in lidar sampling gaps with return intervals on the scale of days [12]. For example, Lang *et al*. [9] utilized spaceborne lidar from GEDI (Global Ecosystems Dynamics Investigations) and Sentinel-2 imagery to produce 10-m global canopy height models utilizing bagged regression trees. Additionally, Tolan *et al*. [10] combined GEDI data with sub-meter satellite imagery to predict a sub-meter canopy height model with a self-supervised vision-transformer and convolutional decoder. These products are useful for biomass estimates in large regions, but the coarse resolution of smoothed outputs cannot be meaningfully translated to forest metrics [8]. Consequently, there is a need for tools that can produce high-resolution CHMs with extractable 3D tree metrics.

SatCHM (Satellite Canopy Height Model) utilizes high-resolution satellite imagery to produce canopy height models with discernible tree crowns. This technique was previously introduced by Abolt *et al*. [13]. This work aims to open source an improved version of the code. SatCHM uses MS-net from Santos *et al*. [14], a workflow running on a single GPU for image-to-image regression in which an ensemble of fully convolutional networks. They operate on copies of an input image that have been coarsened, and the ensemble cooperates to yield predictions of the target CHM.

The growing accessibility of GPUs and open-source deep learning libraries has made it computationally practical to train increasingly complex neural networks [15]. Thus, SatCHM was developed to be an accessible tool for researchers, allowing for site-specific and temporally flexible predictions of canopy height.

## Methods

The inputs to SatCHM include sub-meter resolution panchromatic satellite imagery, metadata regarding solar angle and the sensor viewing angle at the time of acquisition, lidar-produced canopy height models, and the underlying topography of the forested region from a digital elevation model (DEM) as shown in Figure 1.

**FIG. 1.**
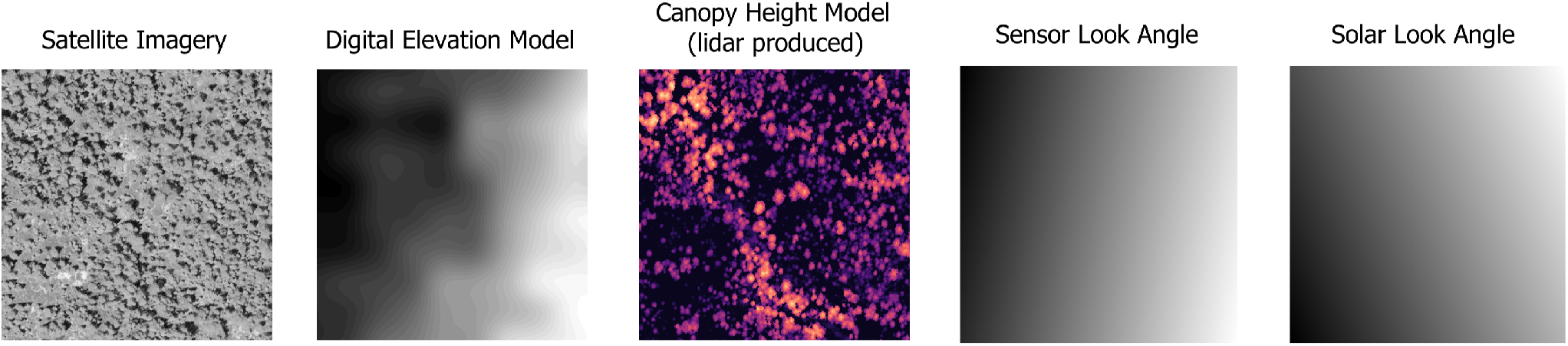
SatCHM inputs. Satellite imagery must be cloudless, panchromatic, and at least sub-meter resolution. Target azimuth, off-nadir angle, solar azimuth, and solar elevation metadata must be documented for each satellite image. Angle metadata generates sensor and solar look angles. Digital elevation models must be at least 30 meters resolution. Lidar-produced canopy height models must be at 0.5 - 0.6 meter or finer resolutions.

Predicted canopy height models underestimate the height of taller canopies and overestimate the height of shorter canopies [8]. Abolt *et al*. [13] discussed the effects of seasonality and solar angle of satellite imagery on prediction accuracy. Model transferability to other forest types is limited by species composition in training data. Therefore, researchers must conduct analyses in context of these error bounds and limitations. Publishing this code enables researchers to improve site-specific height prediction accuracy by incorporating climatic and tree allometric variables as inputs into SatCHM [16–18]. Researchers can then use site-specific model predictions to estimate changes in forest structure and productivity over time and across space. In the example data, canopy height predictions captured the pre-fire forest structure (Figure 2), which can be useful for evaluating interactions between burn severity and pre-fire forest structure [19].

**FIG. 2.**
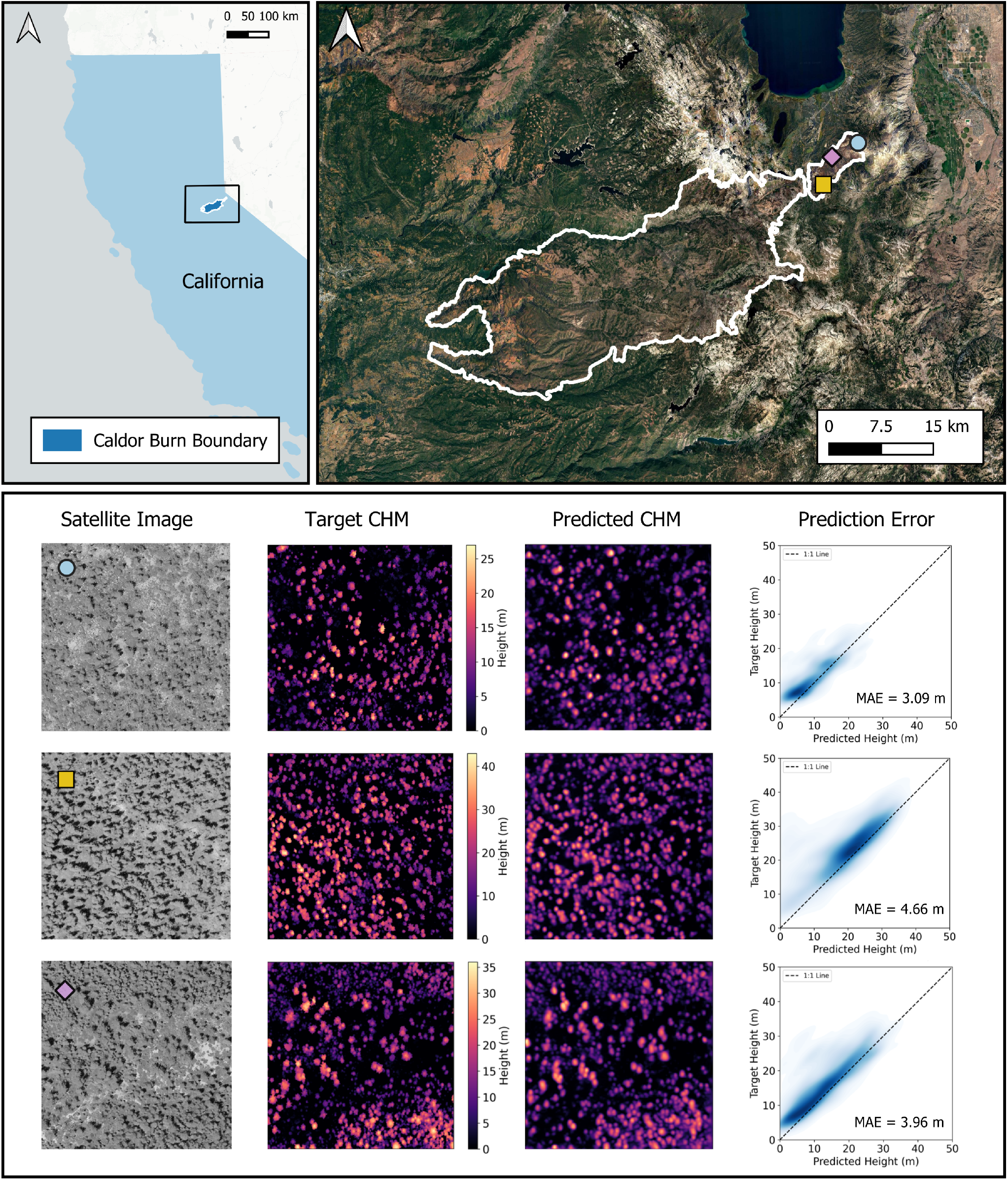
Caldor Fire burn boundary and model predictions. Top left panel shows the Caldor Fire burn boundary in California, USA. Top right panel shows map inset with locations of model predictions. Bottom panel shows the corresponding satellite images, target canopy height models, model-predicted canopy height models, and mean absolute prediction errors.

There are two versions of SatCHM available on the repository. They contain identical neural network architecture and require the same inputs. However, each version is structured for different intended user bases.

Version 0 is intended for a broad set of users interested in forestry and ecological applications of the CHM outputs. Integrated documentation, interactive prompts, and progress indicators are included to enhance usability. The user must manually document solar and sensor geometric data from the satellite image metadata. This includes the target azimuth and off-nadir angles of the satellite and the solar elevation and the solar azimuth specific to the date and time of acquisition of the satellite imagery. Additionally, the user documents the input file paths. SatCHM takes these inputs and tiles them into 512 x 512 pixels (256 x 256 m) for the neural network.

Pixelwise and treewise comparisons are used to evaluate model performance in this version. With pixelwise comparison, each predicted pixel is compared to the corresponding target pixels, but this method suffers from exploding losses and is difficult to interpret. This is because tree crowns are not stable across images and time (satellite images and lidar-produced CHMs); slight spatial shifts in tree positions and variability in crown shapes between predictions and targets can lead to large pixel-wise errors. To address this, treewise comparisons are used as a more interpretable approach. These comparisons rely on a watershed segmentation method to delineate Tree Approximate Objects (TAOs) from the canopy height model [20]. For each TAO, the highest point is identified, and the predicted and target heights are compared using mean absolute error. This provides more useful metrics for understanding and refining predictions, particularly for end-users who are more concerned with tree-level structure than with exact pixel matches [21].

Version 1 refines the previous version by reducing manual script execution and improving software streamlining. Change-log details are documented in the software README.md. This version is designed for users constructing 3D wildland fuels maps. The code is structured in a modular format with main and utils files to ease further engineering efforts and improve interpretability. Its design prioritizes ease of use for the user, minimizing setup effort, manual data downloading, variable specification, and file execution count while retaining all previous functionality from version 0. USGS 3DEP lidar is set to be used as the default lidar source to expand spatial coverage and make training easier, although custom lidar is still supported as in version 0. Treelists are generated with watershed segmentation via cloud2trees to interface cleanly with wildland fuels modeling software, such as QUIC-Fire [21, 22].

The neural network architecture was originally introduced by Santos *et al*. [14] for the prediction of flow fields in porous media. For a comprehensive explanation, readers are encouraged to consult that work. The MS-net architecture consists of branches (“scales”) of convolutional layers operating on different resolutions of the input images. For this application, there are three fully convolutional networks with 15 layers, operating at three different scales coarsened by a factor of 4. Scale 0, with 8 filters, captures the full domain size while scale 2, with 128 filters, operates on the coarsest scale. The loss function used in the training of MS-net accounts for the prediction error at each scale. Each scale’s contribution is calculated as the mean squared error between the predicted field *ŷ* and the corresponding coarsened true field *y*, normalized by the variance of the true field, 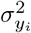. The global loss function is then expressed as a weighted sum of these scale-specific errors [14, 15]. The global loss function *L* is:

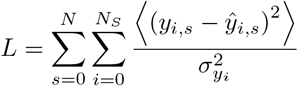

A ReLU activation function was utilized to avoid negative estimates of canopy height [13].

## Results

Example data for demonstration are from a mountain chaparral and mixed conifer forest in eastern Eldorado National Forest, California, USA. Satellite images and lidar-produced canopy height models were collected before the Caldor Fire of 2021 (Figure 2). Satellite images were collected between September 2011 and June 2017. The lidar data was collected in 2010. The preprocessing, training, and validation of 2,037 samples took 2 days using an NVIDIA GeForce RTX 5070 GPU with 32 GB of memory. We used treewise metrics to evaluate model performance. Mean absolute error (MAE) was independently calculated for each prediction. The median of the MAEs was 6.06 meters.

## Discussion

SatCHM uses repeat satellite imagery with a revisit time of days, enabling high-temporal resolution monitoring of forest structural change [13]. By open-sourcing the code, this work allows researchers to explore other site-specific parameters that can improve height predictions. Several applications of this software include evaluating changes to biomass over time and modeling wildland fuels [3, 19, 21]. Future improvements to this software include building a pipeline to commercially available PLANET imagery.

## Acknowledgments

This study was funded and supported by the Laboratory Directed Research and Development under ‘Experiemental Research’ at Los Alamos National Laboratory. The authors thank the FIRE team and others in the Earth and Environmental Sciences division, including but not limited to Julia Oliveto, Javier Santos, and Rod Linn.

